# Modelizing *Drosophila melanogaster* longevity curves using a new discontinuous 2-Phases of Aging model

**DOI:** 10.1101/025411

**Authors:** Hervé Tricoire, Michael Rera

## Abstract

Aging is commonly described as being a continuous process affecting progressively organisms as time passes. This process results in a progressive decrease in individuals fitness through a wide range of both organismal – decreased motor activity, fertility, resistance to stress – and molecular phenotypes – decreased protein and energy homeostasis, impairment of insulin signaling. In the past 20 years, numerous genes have been identified as playing a major role in the aging process, yet little is known about the events leading to that loss of fitness. We recently described an event characterized by a dramatic increase of intestinal permeability to a blue food dye in aging flies committed to die within a few days. Importantly, flies showing this so called ‘Smurf’ phenotype are the only ones, among a population, to show various age-related changes and exhibit a high-risk of impending death whatever their chronological age. Thus, these observations suggest that instead of being one continuous phenomenon, aging may be a discontinuous process well described by at least two distinguishable phases. In this paper we addressed this hypothesis by implementing a new 2-Phases of Aging mathematiCal model (2PAC model) to simulate longevity curves based on the simple hypothesis of two consecutive phases of lifetime presenting different properties. We first present a unique equation for each phase and discuss the biological significance of the 3 associated parameters. Then we evaluate the influence of each parameter on the shape of survival curves. Overall, this new mathematical model, based on simple biological observations, is able to reproduce many experimental longevity curves, supporting the existence of 2-phases of aging exhibiting specific properties and separated by a dramatic transition that remains to be characterized. Moreover, it indicates that Smurf survival can be approximated by one single constant parameter for a broad range of genotypes that we have tested under our environmental conditions.

**Author Summary:** The perception we can have of a process directly affects the way we study it. In the literature, aging is generally described as being a continuous process, progressively affecting organisms through a broad range of molecular and physiological changes ultimately leading to a dramatic decrease of individuals’ life expectancy. As such, aging studies focus on changes occurring in groups of individuals through time, considering individuals taken at a given time as being all equivalent. Instead, the recently described Smurf phenotype [1] suggested that any given time, a population could be divided in two subpopulations each characterized by a significantly different risk of impending death.

By formalizing here the concept of a discontinuous aging process using a mathematical model based on simple experimental observations, we propose a theoretical framework in which aging is actually separated in two consecutive phases characterized by three parameters easily quantifiable *in vivo*. Thus, the model we present here brings new tools to assess the events occurring during aging using a novel angle that we hope will open a better understanding of the very processes driving aging.

## Introduction

Although considerable progress has been made towards the identification of genetic factors influencing longevity, numerous fundamental questions remain about aging, including the nature of the aging process and the ways aging leads to organismal death. Works based on model organisms such as the nematode *Caenorhabditis elegans* and the fly *Drosophila melanogaster* have allowed the identification of genes and signaling pathways that play an evolutionarily conserved role in the modulation of longevity. Those genes are involved in various processes such as immunity, protein homeostasis, energy homeostasis, stress resistance or tissue homeostasis maintenance. Many theories have been proposed to tie up the diversity of observations into a model that would explain the involvement of those various processes in the aging phenomenon. Ranging from the oxidative stress theory of aging [2] to the resource allocation theory [3] through pleiotropic antagonism [4], the proposed theories tend to highlight the apparent balance taking place in an organism between maintaining the individual alive and maximizing the probability of maintaining the population (or the species) across time and environmental variations. A common feature of all these theories is that they rely on progressive continuous changes of one or some parameters along lifespan that sustain an age-dependent exponentially increasing mortality rate. For instance, increased levels of inflammation, impairment of insulin signaling, decreased energy stores are commonly accepted as being hallmarks of aging that progressively evolve with chronological age [5]. This view of aging as being a continuous process has been popularized since the birth of the aging field, as illustrated by Pearl’s rate of living theory [6]. However recent data in several organisms suggest that the way to death may be paved with non-continuous events that allow to discriminate between several distinct populations at a given chronological age – the absolute time since birth.

In *D. melanogaster*, such a dramatic transition occurring in individual flies prior to death was recently described [1]. By feeding flies using a food dye that is normally not absorbed by the drosophila digestive tract, we could identify, at different chronological ages, individuals showing an extended blue coloration where most of the flies showed a blue color restricted to the proboscis and digestive tract (Fig 1A). We showed that the proportion of individuals characterized by this phenotype increases quasi linearly as the population ages. Further characterization of these individuals, that we named Smurfs because of their blue coloration, allowed us to identify a set of co-segregating phenotypes. Whatever their chronological age, compared to their non-Smurfs counterpart, Smurfs show many hallmarks of aging, such as a significantly increased expression of antimicrobial peptides (AMPs), increased expression of FOXO targets, decreased energy stores (glycogen and triglycerides – TG), decreased spontaneous motor activity and a dramatic increase in the probability of death. Strikingly, Smurfs show this set of co-segregating characteristics whatever their chronological age although these changes were negligible in age-matched non-Smurfs individuals. Therefore, the continuous modifications in aging hallmarks observed at a population level must be reinterpreted as occurring from the evolution of the Smurfs/non-Smurfs ratio along lifespan. In addition, we showed that all individuals died as Smurfs, indicating that every individual undergoes the phase 1 (non-Smurfs)/phase 2 (Smurfs) transition prior to death.

**Figure 1:**
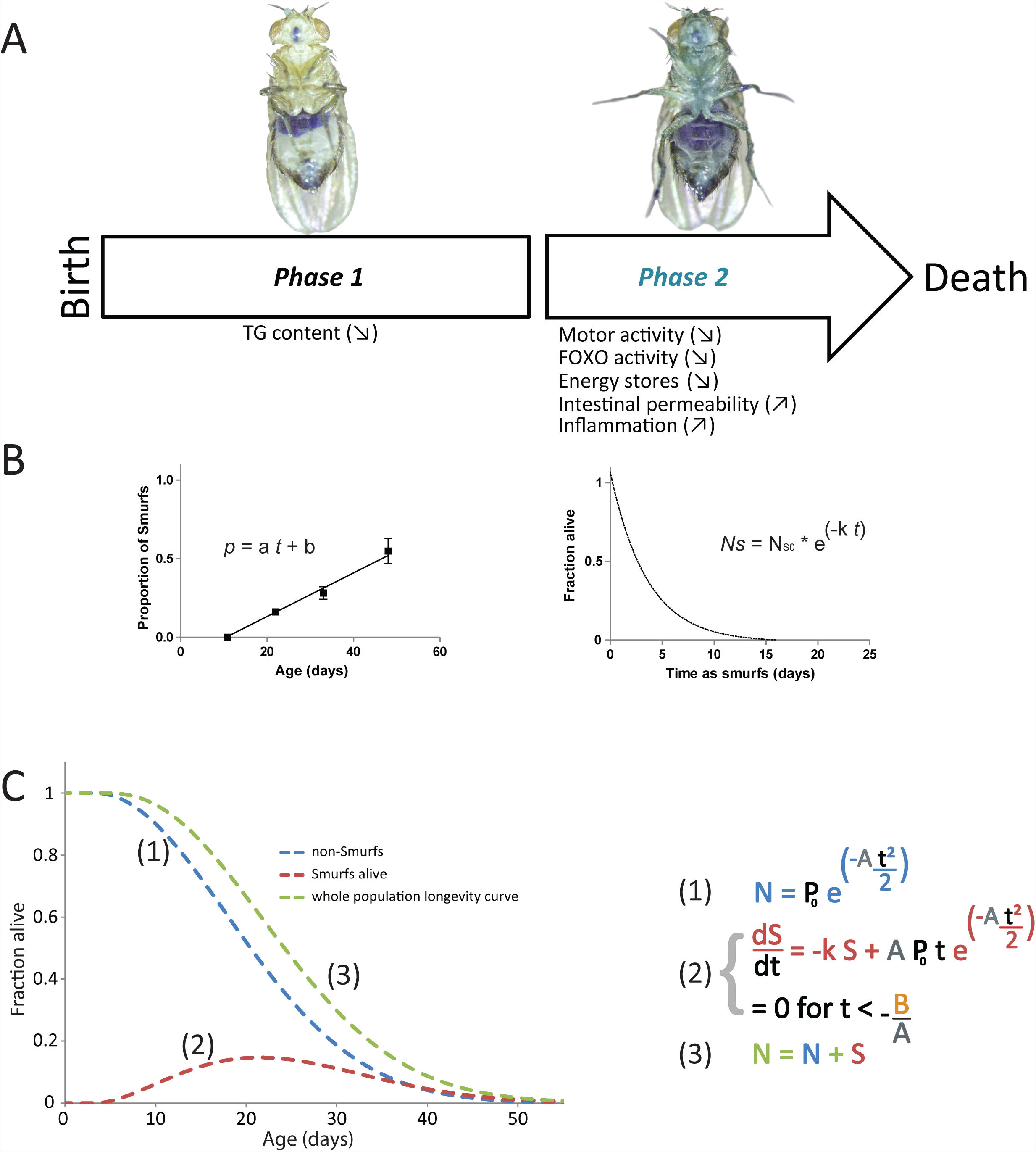
Aging is a 2-phases process. **A.** Aging is characterized by two distinct and consecutive phases. <u>Phase 1</u> is characterized by a time-dependent increase in the probability of at least one organ – the intestine – to fail. <u>Phase 2</u> is the terminal phase of life during which a large number of so-called age-related phenotypes occur concomitantly. **B.** Each phase can be described by a distinct equation. Phase 1 is defined by a linear equation (***y* = a *t* + *b*** – left panel) describing the time-dependent increase of the probability for an individual to turn Smurf. Phase 2 is characterized by a 1-phase exponential decay equation (***y = e^-kt^***) – right panel) describing the survival of the Smurf subpopulation at a given chronological time. **C.** The longevity curve of a homogenous population (green line) of flies is the sum of the number of non-Smurfs flies (blue line) and living Smurfs (red line). The mathematical equations of the different curves are given in the right panel. The model uses 3 parameters; a is the rate of apparition of the Smurfs in the whole population, ***-b/a*** is the age at which the Smurfs appear in the population and k is the rate constant defining the Smurf longevity.

Existence of a sharp transition prior to death has also been observed recently in worms [7] and may underlie the recent observation of specific metabolic markers that are predictors of death in humans [8,9,10]. Taken together, these data suggest that aging can be separated in at least two distinct phases as described in F 1A that illustrates the Smurf phenotype in flies. In this new perspective of a discontinuous aging process, a theoretical description of aging, amenable to experimental validations or refutations, would be highly beneficial.

In this article, we present a first step towards this goal by implementing a simple theoretical model, the 2PAC model, assuming that aging can be separated in two distinct phases, each one characterized by specific features reflected in mathematical equations. In the first phase of their life individuals benefit from a null mortality rate but show a time-dependent increase of the probability to undergo an abrupt transition towards phase 2 where mortality rate is high. After derivation of equations based on these simple biological assumptions, we show that this model is able to reproduce to a large extent experimental data for several genotypes exhibiting significantly different lifespans. In addition we confirmed experimentally that the life expectancy – defined as the T_50_ – of flies in phase 2 is highly similar across the seven genotypes analyzed, as predicted by the model analysis. This theoretical analysis highlights the interest of re-interpreting longevity experiments by taking into account distinct phases separated by abrupt transitions and raises the question of evolutionary conservation of the events leading to death.

## Results

### A 2-phases model of aging: hypothesis, mathematical description and biological relevance of the different parameters

Our previous data suggest that aging can be separated in two distinct phases as described in Fig 1A. Thus, at any time point, a total number of individuals N_T_ will be shared between N individuals in phase 1 (“non-Smurfs”) and S individuals in phase 2 (“Smurfs”). Based on experimental observations (notably the constant median lifespan of individuals in phase 2 whatever their chronological age), we here propose that the evolution of these two populations present in these different phases can be described mathematically by simple coupled equations. First, individuals in phase 1 are considered as exhibiting a null mortality rate and transition to phase 2 is an essential prerequisite to death, in agreement with experimental data presented in [1] as well as in the present article (Fig 3). However we assume that they have a probability **p** to become Smurfs when their age exceeds a threshold **t_0_=-b/a** and that this probability increases linearly as a function of time, in agreement with our previous observations (Fig 1B *left panel*): **p**= ***a*** × *t* + ***b***. In phase 2, we assume that individuals have a constant probability of death per unit of time ***k***, so that an isolated population of **Ns**_**0**_ Smurfs individuals follows a one-step exponential decay equation ***Ns = Ns_**0**_ × e^-k×t^*** (Fig 1B *right panel*).

**Figure 3:**
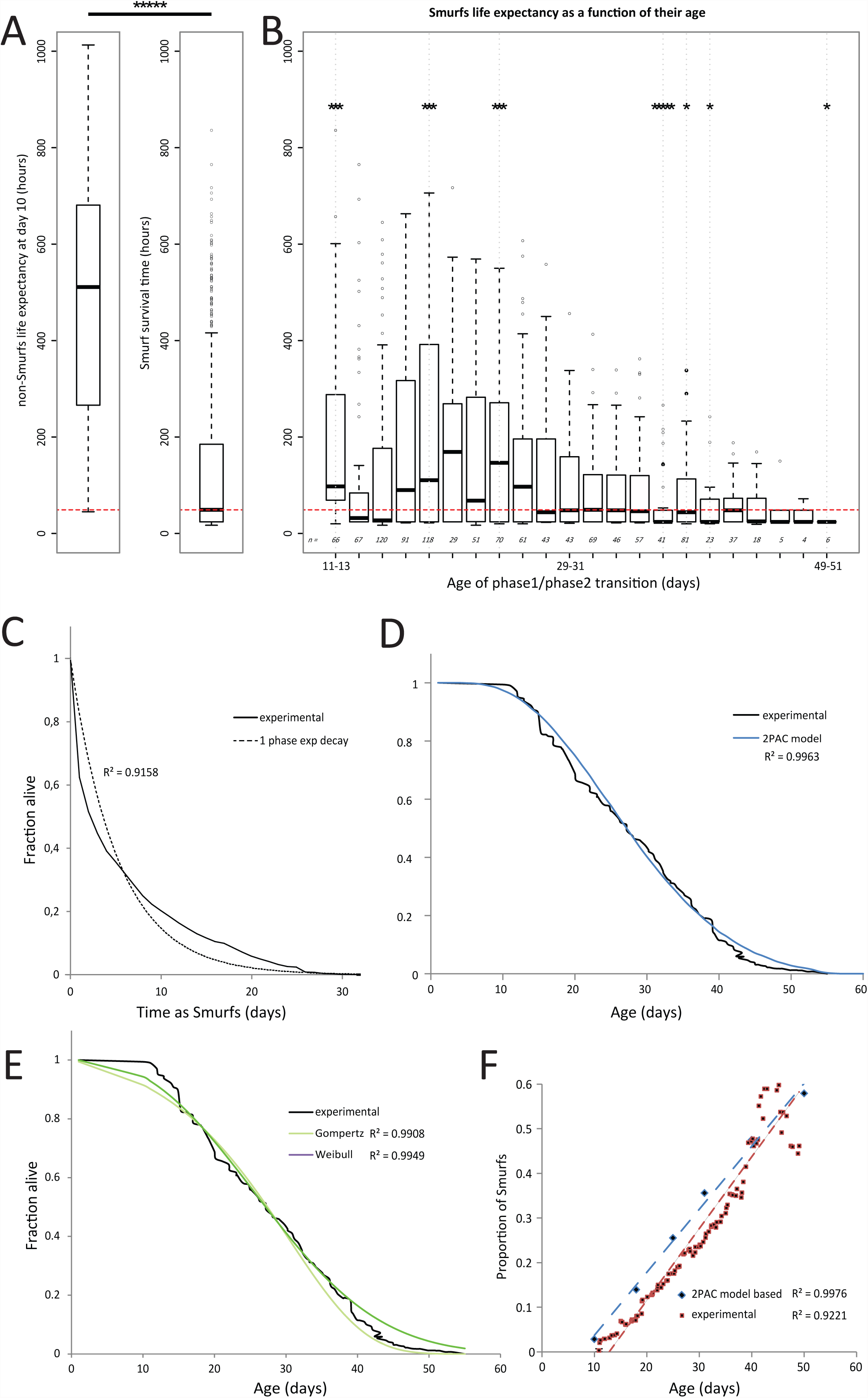
Smurf death rate can be considered as chronological-age independent in *drsGFP* females. **A.** Median life expectancy of 10 days old females (*left panel*, 21.29 days) is significantly different the median survival time of Smurfs (*right panel*, 2.04 days) (*****, p < 10^-5^). **B.** The majority of Smurfs grouped by 48 hours (746 out of 1146 individuals) shows a median ‘survival time as Smurfs’ that is not significantly different from the ‘Smurf survival time’ calculated using the whole Smurf population (p > 0.05, no *). Thus we will use this distribution to generate an average ‘Smurf survival curve’. **C.** Survival curves of Smurf flies from a population of mated *drsGFP* females monitored daily for their Smurf status and death. The equation of that average ‘Smurf survival curve’ was then determined using non-linear regression based on a 1-phase exponential equation ***e***^***-kt***^ with ***k*** = 0.1911 (IC [0.1694 to 0.2129]) R^2^ = 0.9158. **D-E.** The 2PAC model allows to fit the experimental longevity curve with a precision (***a***_2PAC_ = 0.0039; ***b***_2PAC_ = -0.019; R^2^ = 0.9963) similar to the fits obtained with either the Gompertz model (***A***_Gompertz_ = 0.0053; ***k***_Gompertz_ = 0.0942, R^2^ = 0.9908) or the Weibull model (***a*** Weibull = 0.000485; ***k***_Weibull_ = 2.4746; R^2^ = 0.9949). **F.** Comparison of the experimental (0.01607 ± 0.0004693; R^2^ = 0.9221) and theoretical (0.01512 ± 0.0003713; R^2^ = 0.9976) SIRs. The goodness of fit was calculated with both Pearson (p < 0.0001) and Spearman tests (p < 0.005). The theoretical SIR calculated with the 2PAC model adjusted parameters is not significantly different from the experimental one (p = 0.5578).

Initially only the non-Smurf population **N** is present. Thus, during the period [0, t_0_] we have N = P_0_ and S = 0, where **P_0_** is the original population.

If we make a change of variable by taking **t_0_** as the origin, we have now a set of two coupled equations:

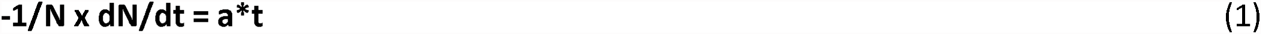

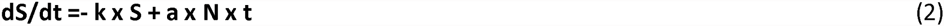

with the initial conditions **N(0)** = **P_0_** and **S(0) = 0**.

These equations can be solved analytically (see Fig S1A) but, due to the complex structure of the final equation, calculation may be unstable on standard 64 bits computer for some parameter values thus leading to a chaotic behavior of **N**_**T**_ = **f(t)**. Therefore, as an alternative, the model was implemented in an Excel file (available on request) as an iterative model devoid of instability (*see material and methods*). The resulting curves are of sigmoidal shape as illustrated in Fig 1C.

By separating aging in 2 distinct phases each defined by a specific equation, the Smurf model allows simple biological interpretation of its parameters ***a***, ***b*** and ***k***, more easily than the classical Gompertz [11] and Weibull [12,13] models do. First, the linear phase 1 parameters ***a*** and ***b*** characterize phase 1 properties and are respectively the slope and the *y-axis* interception point of the linear curve describing the probability of phase transition. ***a*** is expressed in ‘additional fraction of Smurfs per unit of time’ and corresponds to the rate of apparition of Smurfs in the population; we will thus name it daily failure rate. For a given ***a, b*** determines the first day Smurfs can be observed in the population. This day is defined for every 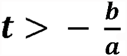. It characterizes the *tolerance* of the population to undergo a phase 1/phase 2 transition. Finally, the ***k*** parameter is the unique parameter defining the rate at which Smurfs die. Since Smurfs are the only individuals dying in the population we will name it *death rate constant*. In the next section we investigate how each of these three parameters affects the final shape of survival curves.

### Influence of the different parameters on the shape of aging curves

To study the effect of the different parameters on the longevity curves (Fig 2), we set a series of initial parameters and then modified these parameters one by one.

**Figure 2:**
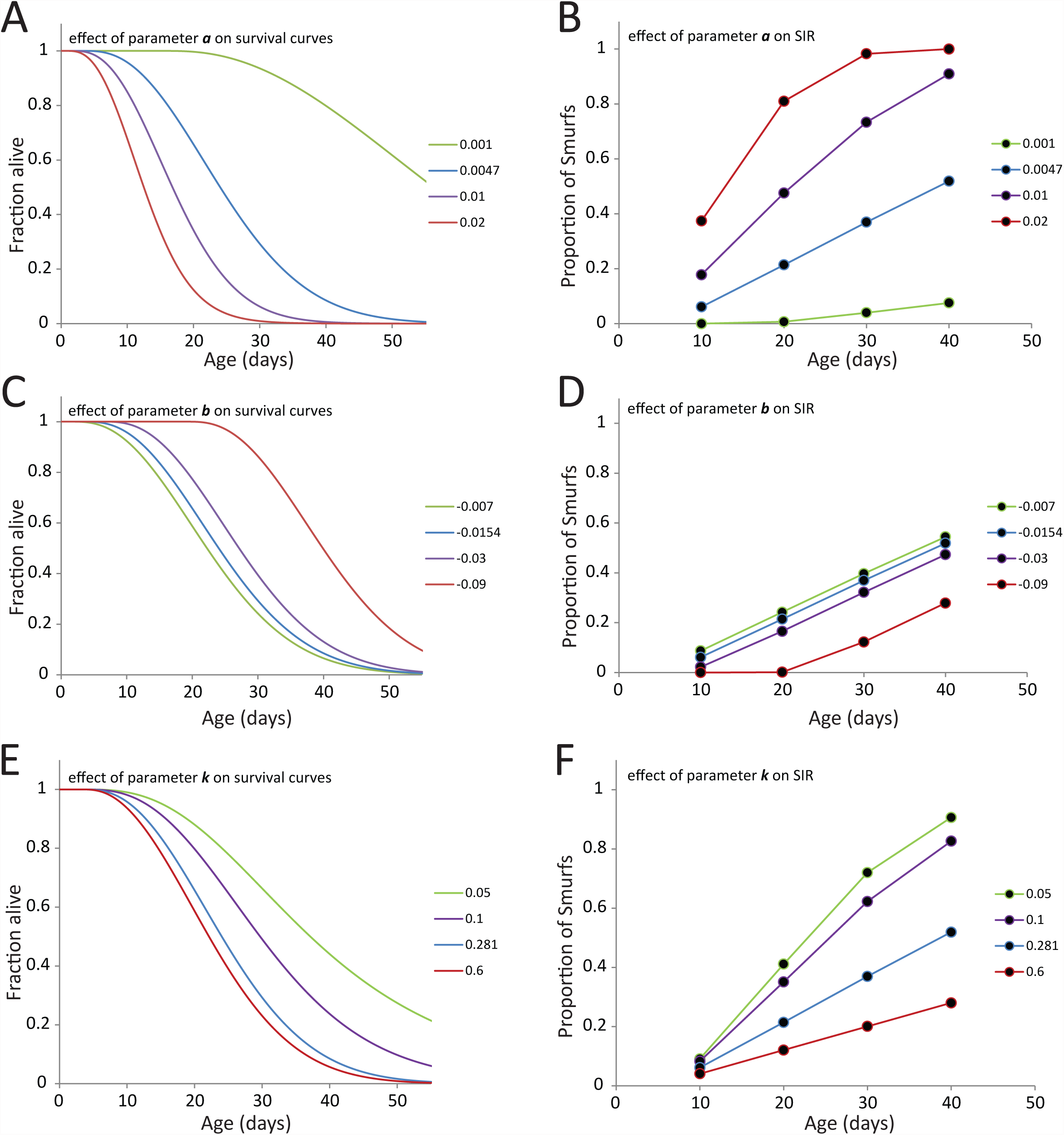
Effects of the different parameters of the model on lifespan. **A, B.** As ***a*** increases, lifespan decreases and Smurf Increase Rate (SIR) increases. ***C, D.*** When ***b*** increases, lifespan increases without affecting the SIR but the first Smurfs appear later. **E, F.** An increase of ***k*** decreases both lifespan and the SIR. Thus, by measuring lifespan and SIR of flies in two distinct conditions indicates which parameter is affected by the treatment.

First, increasing the daily *failure rate* ***a*** results in a dramatic decrease in lifespan of the population, by affecting the length of the initial mortality plateau, the median lifespan T_50_ and the maximum lifespan of the population (Fig 2A). The Smurf Increase Rate (SIR), which is the evolution of the proportion of Smurfs in the population **S**_**t**_**/(S_t_+N_t_)**, also increases with ***b*** (Fig 2B). Secondly, increasing the *tolerance* parameter ***b*** (negative value) increases the length of the initial mortality plateau, the T_50_ and the maximum lifespan of the population (Fig 2C). It also delays the apparition of Smurfs in the population without affecting the rate at which they appear (Fig 2D). It is essential to notice that we only considered negative value for ***b*** in these simulations. For a null or a positive value, the proportion of Smurfs would increase from day 0, thus dramatically decreasing or even suppressing the initial mortality plateau. Finally, increasing the death rate constant ***k*** does not dramatically affect the length of the initial plateau but decreases both the T_50_ and maximum lifespan of the population. Interestingly, as it affects the turnover of the Smurf population, increasing ***k*** decreases the SIR although it decreases lifespan. This is a case that we have not observed experimentally so far as we only observed a negative correlation between SIR and lifespan.

It is important to highlight a few points provided by this study of the effect of the different parameters. Primarily, not all parameters impact lifespan to a similar extent. A 5-fold change of ***a***, ***b*** or ***k*** generates more than a 2-fold change of the T_50_ in the case of ***a*** (Fig 2A, blue versus green curve) that is reduced to a 32% change for ***b*** (Fig 2C, green versus purple curve) and 28% for ***k*** (Fig 2E, blue versus green curve). Then, as described in the precedent paragraph, an increase of the different parameters affects the lifespan and SIR in different ways: ***a*** decreases the lifespan and increases the SIR, where ***b*** decreases the lifespan without affecting the slope of the SIR, and ***k*** decreases both the lifespan and the SIR. Finally, classical study of population-based mortality rates estimates the *‘apparent mortality’* 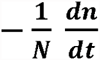 on the whole population. As represented in Fig S1B-D, although only one of the parameters ***k*** affects the mortality *per se* in a population, all the parameters affect the *‘apparent mortality’* calculated at the population level.

### Experimental survival curves can be accurately fitted by the 2-phases aging model

To test whether it is possible to describe experimental longevity curves using this model, we first analyzed the survival of different Smurf populations to determine the best parameter ***k*** describing the exponential decay of these individuals. To do so, we reared a population of 1146 synchronized *drsGFP* [14] mated female flies maintained individually in vials containing blue medium (see material and methods) and each fly was scored for *Smurfness* – whether individuals are Smurf or not – or death daily. We first confirmed that maintaining individual flies throughout life on blue medium did not affect lifespan by comparing their survival curve to one obtained using flies of the same genotype maintained on standard medium by groups of 30 individuals (Fig S2A). Secondly, as previously described in [1] for *w*^*1118*^ flies, we found that the remaining median lifespan of Smurfs is highly similar across life (T_50_ ≈ 2.04 days). Thus, at day 10 for example, the remaining lifespan of *drsGFP* Smurfs is significantly decreased compared to the life expectancy of 10 days old non-Smurf *drsGFP* female flies (Fig 3A). Consistent with this raw estimation of phase 2 individuals’ remaining lifespan, we found that the remaining lifespan of Smurf individuals obtained at all ages during this assay showed limited differences with lifespan of Smurfs obtained at specific ages (Fig 3B) although we noticed a trend towards decreased lifespan for older Smurf flies as well as a higher lifespan variability for the youngest ones. These findings support our model hypothesis that all phase 2 individuals die at a similar pace, modeled by the ***k*** parameter, whatever their age. We then determined the ***k*** parameters of the Smurfs survival equation using a one phase exponential decay fitting curve (Fig 3C).

With this static value of ***k***, we then obtained the remaining two parameters ***a*** and ***b*** with an iterative fitting procedure working as such: starting with a virtual population of non-Smurfs individuals at t = 0, we calculated, with the two previously described model equations, the proportion of individuals undergoing the phase 1/phase 2 transition for every time point until no individuals remain in the non-Smurfs population. Then, every population of Smurfs generated for **t_0_** to **t**_**final**_ decays with the constant rate ***k***. For each time-point, the number of survivors in the whole population is then the sum of non-Smurfs remaining in the initial population and the number of Smurfs still alive. The resulting simulated longevity curve is then fitted to the experimental longevity curve by adjusting ***a*** and ***b*** (***k*** is kept constant) until a maximum is reached for the R^2^ value. The result of the fitting we obtained is presented in the left panel of Fig 3D (R^2^ = 0.9963). Fitting of the experimental data with the classical Gompertz and Weibull models give similar R^2^ values (Fig 3E). To confirm that the model is consistent with other experimental results we calculated the expected Smurf Increase Rate (SIR), based on the parameters used to fit the experimental longevity curve and compared these theoretical values to the experimental data. We found that the theoretical SIR is not significantly different from the experimental one (Fig 3F). It is thus possible to describe the longevity curve of the *drsGFP* mated female population by using the 2-phases aging model based on the assumptions that every individuals die as Smurfs, at a similar pace whatever their age. We then wanted to test whether Smurfs from populations of different genetic backgrounds showing significantly different lifespans are characterized by the same ***k*** or a genotype-dependent one.

### Phase 2 mortality rates are constant for several genotypes presenting different longevities

We previously showed that the rate at which the proportion of Smurfs increases in a population negatively correlates with the T_50_ of that population [1]. We identified 6 lines – from the Drosophila melanogaster Genetic Reference Panel (DGRP) [15] –, characterized by significantly different lifespans showing T_50_ values ranging from 32 to 57.7 days (Fig 4A). Using the previously described methodology to identify Smurf flies, we isolated Smurfs at different ages (15, 23, 33, 41, 47, 55 and 62 days) from the 6 populations of DGRP flies presented in Fig 4A and monitored their remaining lifespan (Fig 4B). Although these survival curves don’t totally overlap they are highly similar, with T_50_ values showing no significant differences with the *drsGFP* large dataset (Fig 4C).

**Figure 4:**
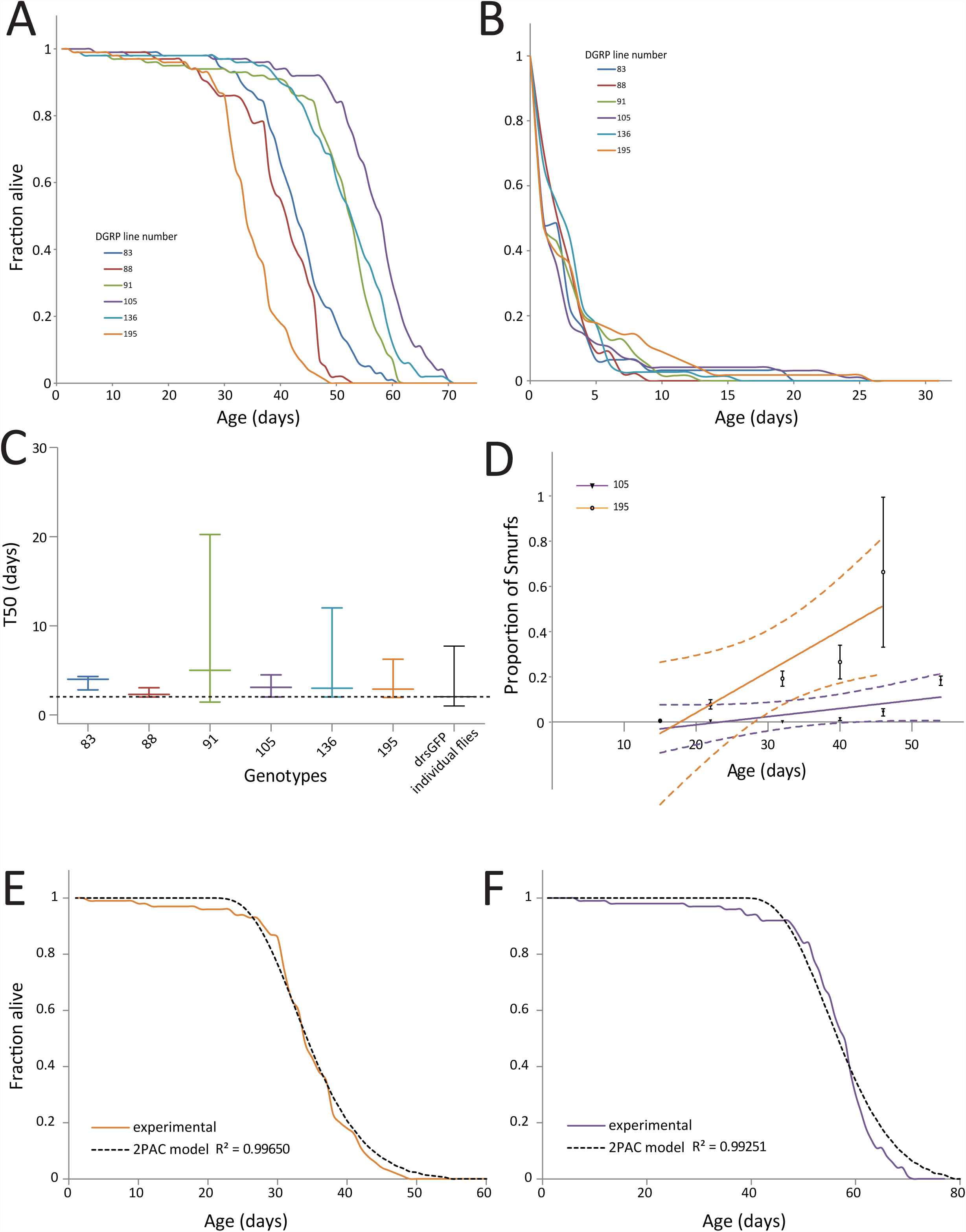
The remaining lifespan of individuals in phase 2 is similar in different drosophila strains. **A.** Mated females from populations of 6 different genetic backgrounds show significant different lifespan curves, DGRP_83 (T_50_ = 42 days; n = 128), DGRP_88 (T_50_ = 39.6 days; n = 127), DGRP_91 (T_50_ = 52.7 days; n = 340), DGRP_105 (T_50_ = 57.1 days; n = 286), DGRP_136 (T_50_ = 53.4 days; n = 243) and DGRP_195 (T_50_ = 32.9 days; n = 262). B, C. The life expectancies of Smurfs from the 6 DGRP lines are highly similar, DGRP_83 (T_50_ = 4.0 days; n = 31), DGRP_88 (T_50_ = 2.3 days; n = 45), DGRP_91 (T_50_ = 5.0 days; n = 96), DGRP_105 (T_50_ = 3.1 days; n = 75), DGRP_136 (T_50_ = 3.0 days; n = 56) and DGRP_195 (T_50_ = 2.9 days; n = 63). In addition, none is different from the one measured using 1146 *drsGFP* individual flies (p > 0.05, 1-way ANOVA using the *drsGFP* as reference) although the Smurf survival measurement protocol was different. Error bars represent median ± s.e.m. D-F. Although SIRs of DGRP_195 (0.01832 ± 0.001602; R^2^ = 0.5612) and DGRP_105 (0.003623 ± 0.001602; R^2^ = 0.8127) are significantly different (p = 0.01579, N > 5 vials per genotype), it is possible to model the longevity curves of the two genotypes using the same ***k*** (phase 2) parameter (calculated from *drsGFP* Smurf flies – figure 3C) with R^2^ > 0.99. Error bars represent mean ± s.e.m. *<u>Note concerning figure 4B and C:</u> the T*_50_ *are higher in figure C than B and this is due to averaging individual vials for the ANOVA test instead of calculating one T*_50_ *using the whole population.*

Thus, under our experimental conditions, the remaining life expectancy (T_50_) of flies in the phase 2 of life, or Smurfs, is almost constant whatever their genotype or the age at which flies underwent the phase 1/phase 2 transition and significantly decreased compared to the non-Smurf flies of the same genotypes.

If this last assumption is acceptable, we expect that it would be possible to reproduce accurately with our model the longevity curves of two populations showing significantly different lifespans. To test this prediction, we decided to fit the longevity curves of two DGRP lines showing significantly different lifespans (DGRP_195, T_50_ = 32.2 days, N = 262 and DGRP_105, T_50_ = 57.7 days, N = 286) as well as significantly different SIRs (Fig 4D) using the Smurfs ***k*** parameter determined in Fig 3C using the *drsGFP* population (T_50_ = 28.8 days, N = 1146). We obtained high quality fits of the experimental longevity curve with a R^2^ > 0.992 for all the two genotypes (Fig 4E and F), a fitting quality similar to those obtained with the Gompertz and Weibull models (Fig S2B and C). We calculated the model-based SIRs for each genotype and compared it to the experimentally determined one. Although the model tends to slightly overestimate the SIR, no statistically significant differences could be found (Fig S2D and E) (p > 0.5). More importantly, as for their experimental counterparts, the theoretical SIRs of the DGRP_195 and DGRP_105 populations are different (p = 0.0013). Thus data derived from the model are compatible with the hypothesis that Smurfs of different genotypes die at a similar pace.

Taken together these results suggest that the Smurf phase, or phase 2 of aging, is highly stereotyped, first on the biochemical aspect as we previously showed [1], but also in respect of the survival of individuals that have underwent the phase 1/phase 2 transition. Moreover, the duration of this last phase of life shows limited differences whatever the chronological age or genotype of the flies under our experimental conditions. This last assumption of the mathematical model is strongly supported by the experimental data obtained with Smurfs survival curves.

## Discussion

In species showing gradual senescence such as *Homo sapiens* and *Drosophila melanogaster*, it is widely accepted that aging manifests itself by a progressive age-dependent decline of fitness accompanied by progressive alterations of biological functions and specific molecular signatures. For example, a conserved signature for age related transcriptome modifications in drosophila and other species is a progressive increase in the expression level of inflammation markers such as anti-microbial peptides (AMPs) in fly or pro-inflammatory cytokines in mammals. In contrast to this view, we propose that individuals issued from a synchronized aging population undergo sharp transitions between states presenting different properties. In this paper, we present a modelization of this assumption in Drosophila, where experimental data have shown that at any time a population can be divided in at least 2 types of individuals, the non-Smurfs and the Smurfs, based on their intestinal permeability.

The new mathematical model of aging that we propose, the 2-PAC model, describes the probability of transitions between these two states as well as the evolution of the different populations in each state. We show that it could accurately reproduce experimental longevity curves from various genotypes over a wide range of median lifespan. This model also brings new and clearer biological interpretation of model parameters than previous parametric models aimed to describe longevity curves. By separating aging in 2 distinct phases, each characterized by a single equation, we could isolate 3 parameters ruling longevity curves that are easily interpretable. *The daily failure rate* ***a*** parameter describes the rate at which individuals enter the second phase of aging. The *tolerance* ***b*** parameter is, with ***a***, an important determinant for the onset of mortality in the population, since it fixes the time **t**_**0**_ **=-*b***/***a*** where the first short living individuals of phase 2 appears in the population. Finally, the death rate constant ***k*** parameter describes the characteristic time constant **τ =1/k** of this phase 2 population lifespan. Surprisingly, ***k*** was found to be mostly constant across lifespan for Smurfs individuals of a given genotype collected every ten days [1]. Here we expanded this result, first by performing a longitudinal analysis with improved time resolution and secondly as we showed that ***k*** is mostly constant between individuals of distinct genetic backgrounds characterized by significantly distinct life expectancies. However, this surprising result has been obtained in our own environmental condition characterized notably by food composition and fixed temperature (see material and methods), and we cannot exclude that different treatments affecting lifespan might affect the ***k*** parameter. We suggest that researchers interested in the study of aging using the drosophila model organism, systematically assess the ***a***, ***b*** and ***k*** parameters in their experimental designs modulating lifespan so that it will be possible to generate a large set of data linking environmental and genetic treatments to the corresponding set of model parameters. This should provide new information on the mechanisms that affect these parameters and to what extent these 3 parameters are independent.

Interestingly, our 2PACs model is fully compatible with previous experimental observations. For instance, our model can easily explain short term variations of death rate **(1/N × dN/dt)** that have been observed in manipulating food composition [16]. Since mortality in our model is essentially controlled by the percentage of individuals in phase 2, any changes in the ***a*** or ***b*** parameters affecting this proportion will quickly impact the mortality rate. Similarly, if we turn to the molecular signatures of aging such as inflammation related molecules, we noticed that phase 2 individuals (Smurfs) show a strong increase of expression of AMPs while individuals in phase 1 (non-Smurfs) present a low AMPs expression whatever their chronological age [1]. Therefore the progressive increase of AMPs expression at the level of the whole population can be reinterpreted as arising from the progressive increase of the proportion of individuals of phase 2 showing a high level of AMP expression in the population. Indeed, we checked with available experimental data – characterized by high time resolution [17] – that our model can accurately describe such an evolution (Fig S3).

At this point, it should be stressed that the 2PAC model that we implemented here is based on our interpretation of our experimental data highlighting two distinct phases. However, it can be easily extended to more complex models including higher phase numbers. Indeed the large set of phenotypical changes that were previously detected in Smurfs indicates that numerous organs and molecular pathways are showing defects in those individuals. Whether one or several of them are the limiting elements leading to naturally occurring age-related death has still to be determined. Thus we can imagine that several consecutive events characterized by different **t**_0_ and ***k*** parameters may occur during aging, the “Smurf state” being the one we were able to observe so far thanks to our Smurf Assay. Whatever the number of such events that could be identified in the future, the description of aging as a succession of discontinuous phases will still remain valid.

The evolutionary conservation of numerous genes, pathways and treatments involved in aging may suggest that discontinuous phases of aging may be conserved across species. Indeed, such a dramatic transition in a state preceding death has been described recently in *C. elegans* [7]. It would be of great interest to investigate whether disruption of calcium homeostasis observed in these phase 2 worms occurs also in phase 2 flies. In humans, scientific and medical reports on raise of intestinal permeability and age-associated chronic diseases or even death have considerably increased in the past few years [18,19,20,21]. It is currently not known whether these phenotypes are hallmarks of a transition to a phase 2 state associated to increased probability of imminent death. Interestingly, a recent article [10] showed that four biomarkers in the blood of human beings predict whether otherwise healthy people are at short-term risk of dying from heart disease, cancer, and other illnesses. Although this study bears some limitations and the causality between these markers and death are not clear, one interpretation of this finding is that death from various causes can be predicted in humans whatever the age of the individuals with a remaining survival time of about 5 years (T_50_). Interestingly, the ratio of this survival time relatively to the total lifespan of humans, (approximately 0.06) is of the same order of magnitude as the one observed in flies between phase 2 mean lifespan and total lifespan. Many studies are required to know whether this observation is purely coincidental or may reflect deeper similarities in the aging process between invertebrates and mammals.

The identification of separate discontinuous phases in the aging process (at least in invertebrates) raises also new questions. In fly the duration of the second phase and the molecular changes that are involved in that phase seem to be tightly linked together and strongly stereotyped, since, as we have shown in this paper and in a previous report, it affects in a very similar way distinct genetic backgrounds and individuals of different ages. We propose that the highly stereotyped transition between these 2 phases and more importantly the phase 2 itself are programmed. If so, we plan to identify the nature of the program. We hope that the gene and protein expression studies of both Smurfs and non-Smurfs populations across aging that we are currently conducting will bring new insights into the aging process and rule out whether that transition is under the control of a yet to identify set of genes or whether it is a more stochastic response involving genetic networks and variability of gene expression levels.

### Materials and Methods

#### Fly Stocks

The Drosophila Genetic Resource Panel (DGRP) lines 83, 88, 91, 105, 136 and 195 as well as the transgenic line *drsGFP* [14] were used for detection of intestinal barrier defects during aging. We used the latter line to keep some continuity with previous work[1] and allow quick verification of results without the blue #1 dye.

#### Fly Culture and lifespan

Flies were cultured in a humidified, temperature-controlled incubator with a 12h on/off light cycle at 26 °C in vials containing standard cornmeal medium (0.68% agar, 5.1% Springaline^®^ inactive dried yeasts, 4.3% sucrose and 2.9% corn flour; all concentrations given in wt/vol). Adult animals were collected under light CO_2_-induced anesthesia, housed at a density of 27–32 flies per vial, and flipped to fresh vials and scored for death every 2–3 days throughout adult life.

#### Smurf assay

Unless stated otherwise, flies were aged on standard medium until the day of the Smurf assay. Dyed medium was prepared using standard medium with blue dye #1 added at a concentration of 2.5% (w/v). Flies were kept on dyed medium overnight. A fly was counted as a Smurf when dye coloration could be observed outside of the digestive tract. To calculate the Smurf increase rate (SIR), we plotted the average proportion of Smurfs per vial as a function of chronological age and defined the SIR as the slope of the calculated regression line.

#### Equation Solving

Equations presented in Fig S1A were solved using the online tool www.wolframalpha.com

#### Iterative implementation of the 2PAC model

A virtual population of drosophila initially contains **N_0_** individuals with a probability p for individuals to become Smurf p(0) = 0. As ***t*** increases, p(**t**) becomes non-null and **N**_**t**_ **= N**_**t-1**_ - **p(t-0.5) × N**_**t-1**_. The population of Smurfs that appeared at time ***t***, **p(t-0.5) × N**_**t-1**_, then decays following phase 2 equation. For any ***t***, the number of survivors is **N**_**0**_ minus the number of Smurfs that died until ***t***.

#### Statistical analysis

Linear regression lines of Smurf proportion during aging were determined using at least 16 individual points (4 time points and 4 replicates per time point) in GraphPad Prism version 5. Correlation of the datasets was assessed using the Pearson test for linear regressions as implemented in the software. Comparison of slopes and testing for non-null slope values were done using GraphPad Prism. Median lifespans were tested for significant differences using the Wilcoxon test implemented in R version 3.1.2. All statistical tests were two-sided.

## Acknowledgements

We thank Scott D. Pletcher for providing raw transcriptomic data from [17].

## Bibliography

1. Rera M, Clark RI, Walker DW (2012) Intestinal barrier dysfunction links metabolic and inflammatory markers of aging to death in Drosophila. Proc Natl Acad Sci U S A.

2. Harman D (1956) Aging: a theory based on free radical and radiation chemistry. J Gerontol 11: 298–300.

3. Medawar PB (1952) An unsolved problem of biology. Med J Aust 1: 854–855.

4. Williams GC (1957) Pleiotropy, Natural Selection, and the Evolution of Senescence. Evolution 11: 398–411.

5. Lopez-Otin C, Blasco MA, Partridge L, Serrano M, Kroemer G (2013) The hallmarks of aging. Cell 153: 1194–1217.

6. Pearl R (1922) The Biology of Death: J. B. Lippincott Company.

7. Coburn C, Allman E, Mahanti P, Benedetto A, Cabreiro F, et al. (2013) Anthranilate fluorescence marks a calcium-propagated necrotic wave that promotes organismal death in C. elegans. PLoS Biol 11: e1001613.

8. Pinto JM, Wroblewski KE, Kern DW, Schumm LP, McClintock MK (2014) Olfactory dysfunction predicts 5-year mortality in older adults. PLoS One 9: e107541.

9. Horne BD, May HT, Muhlestein JB, Ronnow BS, Lappe DL, et al. (2009) Exceptional mortality prediction by risk scores from common laboratory tests. Am J Med 122: 550–558.

10. Fischer K, Kettunen J, Wurtz P, Haller T, Havulinna AS, et al. (2014) Biomarker profiling by nuclear magnetic resonance spectroscopy for the prediction of all-cause mortality: an observational study of 17,345 persons. PLoS Med 11: e1001606.

11. Gompertz B (1825) On the nature of the function expressive of the law of human mortality, and on a new mode of determining the value of life contingencies. Phil Trans Roy Soc: 513–585.

12. Weibull W (1951) A Statistical Distribution Function of Wide Applicability. ASME Journal of Applied Mechanics: 293–297.

13. Fréchet M (1927) Sur la loi de probabilité de l’écart maximum. Ann Soc Polon Math

14. Ferrandon D, Jung AC, Criqui M, Lemaitre B, Uttenweiler-Joseph S, et al. (1998) A drosomycin-GFP reporter transgene reveals a local immune response in Drosophila that is not dependent on the Toll pathway. Embo J 17: 1217–1227.

15. Mackay TF, Richards S, Stone EA, Barbadilla A, Ayroles JF, et al. (2012) The Drosophila melanogaster Genetic Reference Panel. Nature 482: 173–178.

16. Partridge L, Pletcher SD, Mair W (2005) Dietary restriction, mortality trajectories, risk and damage. Mech Ageing Dev 126: 35–41.

17. Pletcher SD, Macdonald SJ, Marguerie R, Certa U, Stearns SC, et al. (2002) Genome-wide transcript profiles in aging and calorically restricted Drosophila melanogaster. Curr Biol 12: 712–723.

18. Farhadi A, Banan A, Fields J, Keshavarzian A (2003) Intestinal barrier: an interface between health and disease. J Gastroenterol Hepatol 18: 479–497.

19. Fasano A, Shea-Donohue T (2005) Mechanisms of disease: the role of intestinal barrier function in the pathogenesis of gastrointestinal autoimmune diseases. Nat Clin Pract Gastroenterol Hepatol 2: 416–422.

20. Sandek A, Rauchhaus M, Anker SD, von Haehling S (2008) The emerging role of the gut in chronic heart failure. Curr Opin Clin Nutr Metab Care 11: 632–639.

21. Harris CE, Griffiths RD, Freestone N, Billington D, Atherton ST, et al. (1992) Intestinal permeability in the critically ill. Intensive Care Med 18: 38–41.

